# Capsular polysaccharide switching in *Streptococcus suis* modulates host cell interactions and virulence

**DOI:** 10.1101/2020.10.12.336958

**Authors:** Masatoshi Okura, Jean-Philippe Auger, Tomoyuki Shibahara, Guillaume Goyette-Desjardins, Marie-Rose Van Calsteren, Fumito Maruyama, Mikihiko Kawai, Makoto Osaki, Mariela Segura, Marcelo Gottschalk, Daisuke Takamatsu

**Affiliations:** Division of Bacterial and Parasitic Diseases, National Institute of Animal Health, National Agriculture and Food Research Organization, Tsukuba, Ibaraki, Japan; Faculty of Veterinary Medicine, University of Montreal, Saint-Hyacinthe, Quebec, Canada; Division of Pathology and Pathophysiology, National Institute of Animal Health, National Agriculture and Food Research Organization, Tsukuba, Ibaraki, Japan; Department of Veterinary Science, Graduate School of Life and Environmental Sciences, Osaka Prefecture University, Izumisano, Osaka, Japan; Saint-Hyacinthe Research and Development Centre, Agriculture and Agri-Food Canada, Saint-Hyacinthe, Quebec, Canada; Microbial Genomics and Ecology, Office of Industry-Academia-Government and Community Collaboration, Hiroshima University, Hiroshima, Japan; Scientific and Technological Bioresource Nucleus, Universidad de La Frontera, Temuco, Chile; Graduate School of Human and Environmental Studies, Kyoto University, Kyoto, Japan; The United Graduate School of Veterinary Sciences, Gifu University, Gifu, Gifu, Japan

## Abstract

*Streptococcus suis* serotype 2 strains can cause severe infections in both swine and humans. The capsular polysaccharide (CPS) of *S. suis* defines various serotypes based on its composition and structure. Though serotype switching from serotype 2 has been suggested to occur between *S. suis* strains, its impact on pathogenicity and virulence remains unknown. Herein, we experimentally generated *S. suis* serotype-switched mutants from a serotype 2 strain (SS2) that express the serotype 3, 4, 7, 8, 9, or 14 CPS (SS2to3, SS2to4, SS2to7, SS2to8, SS2to9, and SS2to14, respectively). The effects of serotype switching were then investigated with regards to classical properties conferred by presence of the serotype 2 CPS, including adhesion to/invasion of porcine tracheal epithelial cells, resistance to phagocytosis by murine macrophages, killing by murine and porcine whole blood, and dendritic cell-derived pro-inflammatory mediator production. Results demonstrated that these properties on host cell interactions were differentially modulated depending on the switched serotypes. Using a mouse model of systemic infection, SS2to8 was demonstrated to be hyper-virulent, with animals rapidly succumbing to septic shock, whereas SS2to3 and SS2to4 were less virulent than SS2 because of a reduced systemic inflammatory host response. By contrast, switching to serotype 7, 9, or 14 CPSs had little to no effect. Finally, development of clinical signs in a porcine model of infection was only observed following infection with SS2, SS2to7, and SS2to8. Taken together, these findings suggest that serotype switching can differentially modulate *S. suis* host cell interactions and virulence depending on the CPS type expressed.

**Importance:** *Streptococcus suis* serotype 2 is the most frequently type associated with swine and zoonotic infections. While the serotype 2 CPS is required for virulence and pathogenesis, little information is available regarding that of other serotypes and how differences in serotype can directly affect host cell interactions and virulence. Herein, we constructed serotype-switched mutants from a serotype 2 strain and demonstrated that serotype switching can shift and modulate the *S. suis* host cell interactions and virulence *in vivo*. Among the serotype-switched mutants, the mutant expressing the serotype 8 CPS, whose composition and structure are identical to that of the human pathogen *Streptococcus pneumoniae* serotype 19F, was hyper-virulent, whereas mutants expressing the serotype 3 or 4 CPSs had reduced virulence. These results demonstrate that serotype switching can drastically alter *S. suis* phenotype. Consequently, further importance and attention should be given to the phenomenon of serotype switching and the possible emergence of hyper-virulent isolates.

## Introduction

*Streptococcus suis* is an important porcine pathogen and zoonotic agent causing septicemia, meningitis and many other diseases [1–4]. This bacterium has evolutionarily adapted to pigs, with nearly 100% of carriage rate in the upper respiratory tract [4, 5]. *S. suis* strains are serotyped based on structural differences in the capsular polysaccharide (CPS) [2, 4]. Among thirty-five reported serotypes (serotypes 1-34 and 1/2), serotype 2 is responsible for the majority of human clinical cases and is the most frequently isolated from diseased pigs [2]. Serotypes 1/2, 3, 4, 7, 8, 9, and 14 are also frequently isolated from diseased pigs, although their distributions differ depending on the geographic location [2]. Multilocus sequence typing (MLST) for *S. suis* has demonstrated genetic diversity within this species, with more than 1,000 sequence types, and several clonal complexes (CCs) potentially associated with diseases in humans and pigs [2, 6]. Accumulated serotyping and MLST data indicate the presence of different CCs in the population of serotype 2 strains, and several different serotypes in the respective CCs [pubMLST: http://pubmlst.org/ssuis/]. Taken together, this suggests that serotype switching may occur between *S. suis* serotype 2 and different serotype isolates.

The *S. suis* CPS is produced by the repetition of a defined oligosaccharide unit formed by a unique arrangement of various sugars [7]. Indeed, unique CPS structures of serotypes 1, 2, 3, 7, 8, 9, 14, 18, and 1/2 have been previously determined [8–13] (Fig. S1). Furthermore, previous studies have shown that more than 10 genes related to *S. suis* CPS synthesis are clustered on a genomic locus [7, 14]. Alongside, the CPS synthesis gene (*cps* gene) clusters of serotypes 1 and 14 and serotypes 2 and 1/2 are almost identical [7], with their CPS structure differing by the substitution of only a galactose (Gal) for a *N*-acetylgalactosamine (GalNAc) [10] due to a single nucleotide polymorphism in the glycosyltransferase *cpsK* gene [15]. Except for these four serotypes, gene repertoires in the *cps* gene clusters greatly differ between serotypes [7, 14], indicating that up-take of genomic DNA of different serotypes and replacement of *cps* gene cluster by homologous recombination, using flanking sequences of the clusters, is usually required for serotype switching. In *S. suis*, some strains are naturally transformable, with the competent state induced by competence gene products [16, 17]. Although serotype switching in *S. suis* has not yet been demonstrated, these findings suggest that replacement of the *cps* gene clusters may occur in strains in the competent state through up-take of genomic DNA of the other serotype strains from the environment.

Importantly, the serotype 2 CPS has been shown to play critical roles in protection against phagocytosis by innate immune cells and masking of bacterial surface proteins involved in host cell activation [18]. In addition, several studies have demonstrated non-virulence of the isogenic non-encapsulated serotype 2 mutants in murine and porcine models of infection [18]. However, very little information is available regarding the CPS of other *S. suis* serotypes and is restricted to two studies on serotypes 9 and 14 [18, 19]. Furthermore, comparing the virulence of strains from different serotypes is impossible due to the high genotypic variation between strains. Accordingly, it remains unclear whether *S. suis* serotype switching (i.e., differences in CPS structure) can affect host cell interactions and strain virulence, even though serotype switching may occur among *S. suis* strains.

In the present study, serotype-switched *S. suis* mutants were experimentally generated to investigate the impacts of CPS type on the host cell interactions and virulence *in vivo*. The mutants were switched from serotype 2, which is the most important in this species, to serotypes 3, 4, 7, 8, 9, and 14, which are frequently isolated from diseased pigs and found in several CCs with serotype 2 human isolates (CC1, CC20, CC25, CC28, and CC104). Generated mutants have allowed us to study the modulation of the pathogenesis of *S. suis* caused by serotype switching.

## Results

### Generated serotype-switched *S. suis* mutants contain few mutations other than the *cps* locus

Six different serotype-switched mutants (SS2to3, SS2to4, SS2to7, SS2to8, SS2to9, and SS2to14) and non-encapsulated mutant ΔCPS2, from which the *cps* locus was deleted, were generated from the serotype 2 strain P1/7 (hereafter SS2) (Table 1, generated as illustrated in Fig. S2 and Fig. S3). Serotype-switched mutants were confirmed to belong to the correct serotype using classical serological techniques [23].

**Table 1.**
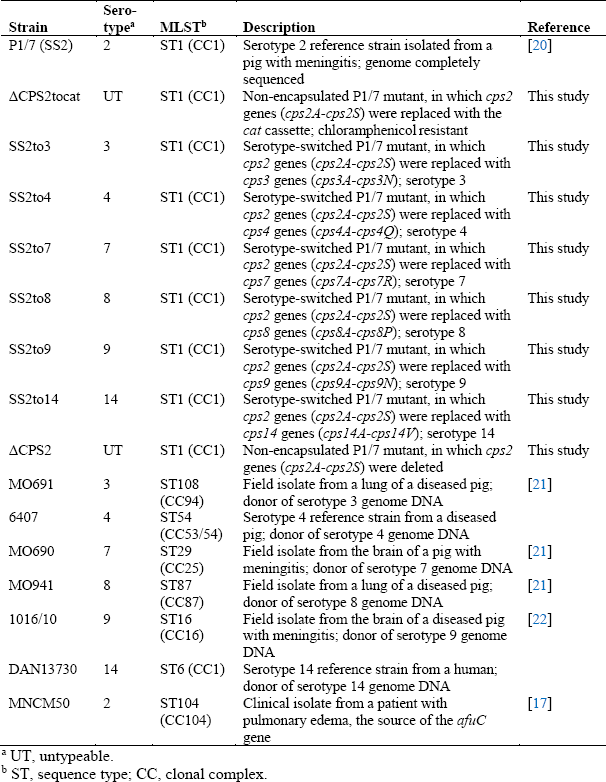
*S. suis* strains used in this study

| Strain | Sero-type<sup>a</sup> | MLST<sup>b</sup> | Description | Reference |
| --- | --- | --- | --- | --- |
| P1/7 (SS2) | 2 | ST1 (CC1) | Serotype 2 reference strain isolated from a pig with meningitis; genome completely sequenced | [20] |
| ΔCPS2tocat | UT | ST1 (CC1) | Non-encapsulated P1/7 mutant, in which *cps2* genes (*cps2A-cps2S*) were replaced with the *cat* cassette; chloramphenicol resistant | This study |
| SS2to3 | 3 | ST1 (CC1) | Serotype-switched P1/7 mutant, in which *cps2* genes (*cps2A-cps2S*) were replaced with *cps3* genes (*cps3A-cps3N*); serotype 3 | This study |
| SS2to4 | 4 | ST1 (CC1) | Serotype-switched P1/7 mutant, in which *cps2* genes (*cps2A-cps2S*) were replaced with *cps4* genes (*cps4A-cps4Q*); serotype 4 | This study |
| SS2to7 | 7 | ST1 (CC1) | Serotype-switched P1/7 mutant, in which *cps2* genes (*cps2A-cps2S*) were replaced with *cps7* genes (*cps7A-cps7R*); serotype 7 | This study |
| SS2to8 | 8 | ST1 (CC1) | Serotype-switched P1/7 mutant, in which *cps2* genes (*cps2A-cps2S*) were replaced with *cps8* genes (*cps8A-cps8P*); serotype 8 | This study |
| SS2to9 | 9 | ST1 (CC1) | Serotype-switched P1/7 mutant, in which *cps2* genes (*cps2A-cps2S*) were replaced with *cps9* genes (*cps9A-cps9N*); serotype 9 | This study |
| SS2to14 | 14 | ST1 (CC1) | Serotype-switched P1/7 mutant, in which *cps2* genes (*cps2A-cps2S*) were replaced with *cps14* genes (*cps14A-cps14V*); serotype 14 | This study |
| ΔCPS2 | UT | ST1 (CC1) | Non-encapsulated P1/7 mutant, in which *cps2* genes (*cps2A-cps2S*) were deleted | This study |
| MO691 | 3 | ST108 (CC94) | Field isolate from a lung of a diseased pig; donor of serotype 3 genome DNA | [21] |
| 6407 | 4 | ST54 (CC53/54) | Serotype 4 reference strain from a diseased pig; donor of serotype 4 genome DNA |  |
| MO690 | 7 | ST29 (CC25) | Field isolate from the brain of a pig with meningitis; donor of serotype 7 genome DNA | [21] |
| MO941 | 8 | ST87 (CC87) | Field isolate from a lung of a diseased pig; donor of serotype 8 genome DNA | [21] |
| 1016/10 | 9 | ST16 (CC16) | Field isolate from the brain of a diseased pig with meningitis; donor of serotype 9 genome DNA | [22] |
| DAN13730 | 14 | ST6 (CC1) | Serotype 14 reference strain from a human; donor of serotype 14 genome DNA |  |
| MNCM50 | 2 | ST104 (CC104) | Clinical isolate from a patient with pulmonary edema, the source of the *afuC* gene | [17] |

<sup>a</sup> UT, untypeable.
<sup>b</sup> ST, sequence type; CC, clonal complex.

Serotype switching had little effect on bacterial growth *in vitro* (Fig. S4). Well-encapsulation of the serotype-switched mutants were confirmed by surface hydrophobicity and transmission electron microscopy (TEM) (Fig. 1A and B). Moreover, purified CPS yields of the mutants SS2to3, SS2to7, SS2to8, SS2to9, and SS2to14 were comparable to those previously reported [9, 11–13] (Table S1). Nuclear magnetic resonance (NMR) analyses confirmed the serotype identity for the serotype-switched mutants, except for SS2to9 (Fig. S5) [9, 11–13]. The CPS of SS2to9 slightly differed from that of serotype 9 strain 1273590 (used for CPS structure determination [11]) in that SS2to9 possessed a glucose instead of a galactose side chain (Fig. S6A), suggesting that the donor strain and SS2to9 may be classified as a serotype 9 variant, which reacts with anti-serotype 9 serum (see Text S1 for more detail). Taken together, these results confirm that the constructed serotype-switched mutants functionally possess and express the CPS of the donor serotype.

**Fig 1.**
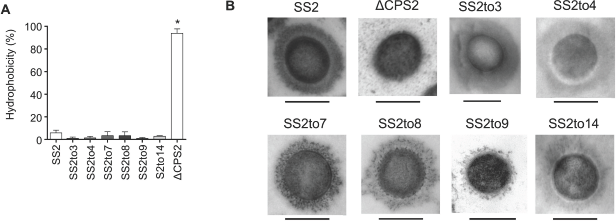
Effect of serotype switching on *S. suis* CPS expression. **(A)** Hydrophobicity of the different *S. suis* strains/mutants. Very low surface hydrophobicity is indicative of high encapsulation, which is demonstrated in the previous study [24]. Data are expressed as mean ± standard error of the mean (SEM) (n = 3). An asterisk denotes a significant difference with SS2 by Mann-Whitney rank sum test (*p* < 0.05). **(B)** Transmission electron micrographs showing CPS expression of the different *S. suis* strains/mutants. Scale bars = 0.5 μm.

To investigate potential mutations in the genomes of the serotype-switched mutants occurred following the transformation of whole genomic DNA, draft genome sequences of the mutants were compared with those of SS2 and the donors. The mutants had mutations in several genes besides the *cps* genes, which differed between mutants (Fig. 2, Fig. S7, and Table S2; see Text S2 for more detail). However, no genes other than *cps* genes were gained in the genomes of the different mutants. Although it remains unclear whether these mutations might affect host-pathogen interactions and virulence, nonsense and frameshift mutations in genes, including virulence-associated genes [18], did not occur (Table S2). This means that the mutants constructed in this study have almost identical genetic background to SS2 compared to the heterogenous genetic background of the different serotype strains, enabling more strict evaluation of the CPS effect hereafter.

**Fig 2.**
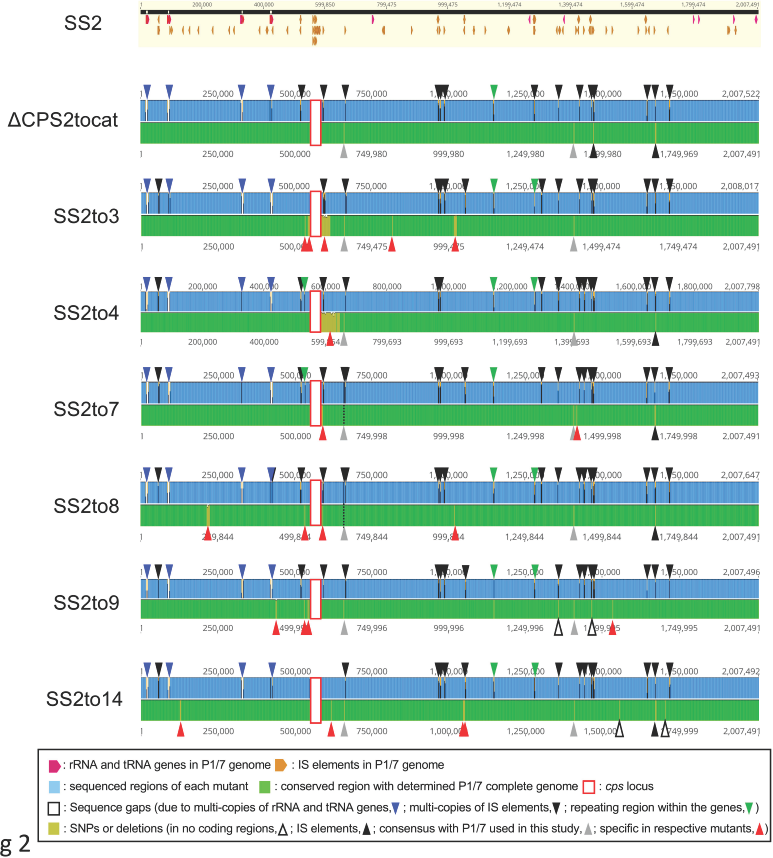
Mutations present in the generated *S. suis* serotype-switched mutants. Each of the schematic representations illustrates the analysis data using Geneious Prime mapping of the draft genome sequence of each mutant (upper part) on the publicly available completed genome sequence of serotype 2 (accession no. AM946016) and the sequence alignment between two genomes (lower part). All gaps between the contigs of each mutant were due to multi-copy genes, such as rRNA genes, tRNA genes and IS elements, or repeated regions within genes. Gaps of the repeated regions within genes were found in the genes corresponding to the SS2 locus tags SSU0496, SSU1127, SSU1171, and SSU1172. Detailed data on mutated genes can be found in Table S2. Below the bottom panel are displayed the descriptions for each color of the different drawings.

### Switching from serotype 2 of *S. suis* can modulate host cell interactions

The serotype 2 CPS has been described to mask surface adhesins involved in the initial interactions with host cells, including adhesion to and invasion of epithelial cells [19, 25], to resist phagocytosis by macrophages and bactericidal killing by blood leukocytes to persist in the bloodstream and cause systemic dissemination [18], and to mask subcapsular immunostimulatory components to interfere pro-inflammatory mediator production by dendritic cells (DCs) [26, 27].

First, using newborn pig trachea (NPTr) cells, the adhesion and invasion capacities were evaluated between SS2 and the mutants. While SS2, SS2to3, SS2to4, SS2to9, and SS2to14 similarly adhered to NPTr cells at 2 h, adhesion of SS2to7 and SS2to8 was significantly greater (*P* < 0.05), similar to that of ΔCPS2 used as a positive control (Fig. 3A). Unlike adhesion results, invasion of the different mutants was similar to that of SS2, with little invasion of NPTr cells overall, although ΔCPS2 showed high levels of invasion, as expected (Fig. 3B).

**Fig 3.**
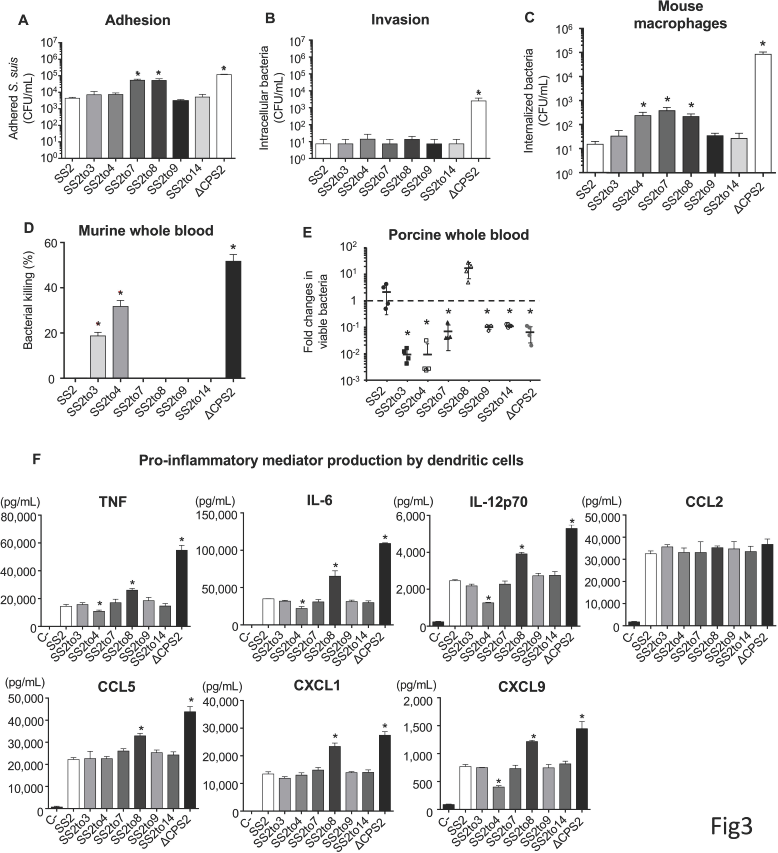
Impact of serotype switching on *S. suis* adhesion to and invasion of porcine tracheal epithelial cells, resistance to phagocytosis by macrophages, whole blood bacterial killing, and pro-inflammatory mediator production by dendritic cells. Adhesion **(A)** and invasion **(B)** of the different *S. suis* strains and mutants to NPTr porcine tracheal epithelial cells after 2 h of incubation. **(C)** Internalization of the different *S. suis* strains and mutants by J774A.1 murine macrophages after 2 h of incubation. **(D)** Killing of the different *S. suis* strains and mutants by murine whole blood after 4 h of incubation. **(E)** Growth capacity of the different *S. suis* strains and mutants in porcine whole blood after 4 h of incubation. **(F)** Pro-inflammatory mediator production by DCs at 16 h following infection with the different *S. suis* strains and mutants as measured by ELISA. Production of tumor necrosis factor (TNF), interleukin (IL)-6, IL-12p70, C-C motif chemokine ligand (CCL) 5, and C-X-C motif chemokine ligand (CXCL) 1, and CXCL9. C-denotes cells in medium alone. All the data represent the mean ± SEM (n = 4). An asterisk denotes a significant difference with SS2 by Mann-Whitney rank sum test **(E)** (*p* < 0.05).

Next, macrophage phagocytosis resistance was evaluated using the J774A.1 murine macrophage cell line. As expected, SS2 and ΔCPS2 were poorly and highly internalized by macrophages, respectively (Fig. 3C). No differences were observed in the internalization between SS2 and the serotype-switched mutants after 1 h incubation (data not shown); however, switching to serotype 4, 7 or 8 significantly increased phagocytosis, after 2 h incubation (*P* < 0.05) (Fig. 3C). However, it should be noted that this increase was of approximately one log-fold, which is, though significant, relatively minor compared to the non-encapsulated mutant (4 log-fold increase).

The capacity to resist the bactericidal effect of leukocytes was then evaluated using murine and porcine whole blood. SS2 was completely resistant to killing by murine blood in contrast to ΔCPS2, which was efficiently killed (60% of killing) (Fig. 3D). While SS2to7, SS2to8, SS2to9, and SS2to14 were also resistant to killing by murine whole blood, SS2to3 and SS2to4 were significantly more killed, with 20% and 30% of killing, respectively (*P* < 0.05) (Fig. 3D). Using a porcine blood system, SS2 was not only able to persist, but also to some extent multiply, whereas ΔCPS2 was markedly cleared (*P* < 0.05) (Fig. 3E). Comparable to SS2, SS2to8 could significantly multiply, whereas all other mutants were cleared at different degrees (Fig. 3E). As with mouse blood, SS2to3 and SS2to4 showed the greatest impairment in their capacity to survive in porcine blood (Fig. 3E). It should be noted, however, that levels of cross-reactive antibodies against the different strains might affect the results observed with the swine blood and thus can be considered a confounding factor, although this fact also mimics the real situation in the field.

Lastly, the interactions with DCs were evaluated. Absence of CPS significantly increased production of all mediators tested (*P* < 0.05), with the exception of CCL2 (Fig. 3F), as previously reported [19, 25]. SS2to3, SS2to7, SS2to9, or SS2to14, along with SS2, did not modulate pro-inflammatory mediator production (Fig. 3F). However, stimulation with SS2to8 significantly increased production of TNF, IL-6, IL-12p70, CCL5, CXCL1, and CXCL9, compared to SS2 (*P* < 0.05) (Fig. 3F). By contrast, SS2to4 induced significantly lower levels of TNF, IL-6, IL-12p70, and CXCL9 than SS2 (*P* < 0.05), but CCL5 or CXCL1. CCL2 production was not modulated regardless of the CPS type (Fig. 3F).

### Serotype switching can differentially modulate *S. suis* virulence in a mouse model of systemic infection

The impact of switching from serotype 2 on *S. suis* virulence was evaluated using a well-established C57BL/6 mouse infection model for *S. suis* serotype 2 virulence studies [28]. Following intraperitoneal inoculation of SS2, 60% of mice died after developing clinical signs of systemic infection (Fig. 4A). By contrast, none of the ΔCPS2-inoculated mice died, presenting no or very mild clinical signs the first 24 h only (Fig. 4A). No significant differences in mortality were observed between SS2 and SS2to3, SS2to7, SS2to9, or SS2to14 (Fig. 4A). However, clinical signs of infection caused by SS2to3 were generally less severe than those by SS2. Unexpectedly, inoculation of SS2to8 significantly increased mouse mortality, with 100% of mice succumbing to septic shock within 24 h post-infection (*P* < 0.05) (Fig. 4A). By contrast, none of the SS2to4-infected mice died, presenting transient clinical signs within the first 48 h (*P* < 0.05) (Fig. 4A).

**Fig 4.**
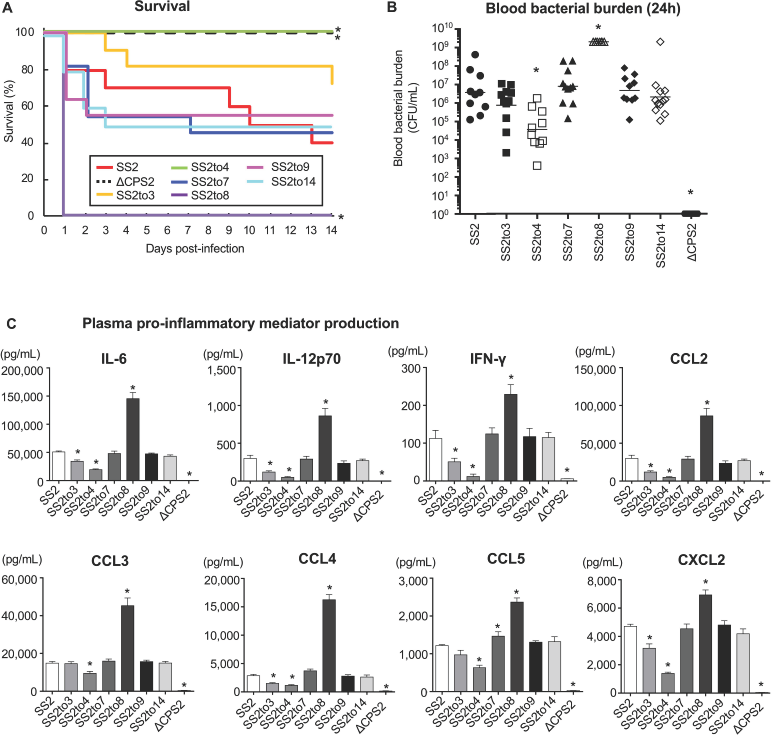
Impact of serotype switching on *S. suis* virulence and plasma pro-inflammatory mediator production in a mouse model of infection. **(A)** Survival of C57BL/6 mice following intraperitoneal inoculation of 1 × 10^7^ CFU of the different *S. suis* strains and mutants. **(B)** Blood bacterial burden 24 h post-infection of C57BL/6 mice. A blood bacterial burden of 2 × 10^9^ CFU/mL, corresponding to average burden upon euthanasia, was attributed to euthanized mice. **(C)** Plasma levels of IL-6, IL-12p70, IFN-γ, CCL2, CCL3, CCL4, CCL5, and CXCL2 in C57BL/6 mice at 12 h following intraperitoneal inoculation of 1 × 10^7^ CFU of the different *S. suis* strains and mutants. Data represent survival curves **(A)** (n = 10-12), geometric mean **(B)** (n = 10-12) or mean ± SEM **(C)** (n = 8). An asterisk denotes a significant difference with SS2 by Log-rank (Mantel-Cox) test **(A)** and Mann-Whitney rank sum test **(B-C)** (*p* < 0.05).

Blood bacterial burdens of infected mice were also determined to investigate the effect on persistent bacteremia. Twenty-four hours post-infection, bacterial burdens of SS2-infected mice averaged 3 × 10^7^ colony-forming unit (CFU)/mL, whereas those in mice infected with ΔCPS2 were not detectable (< 1 × 10^2^ CFU/mL) (Fig. 4B). Similar to mortality, no significant difference was observed between SS2 and SS2to3, SS2to7, SS2to9 or SS2to14 (Fig. 4B and Fig. S8). Meanwhile, blood bacterial burden of SS2to8-infected mice was significantly greater than that of SS2-infected mice (*P* < 0.05), averaging 2 × 10^9^ CFU/mL (Fig. 4B). By contrast, blood bacterial burden was significantly reduced in SS2to4-infected mice compared to SS2 (*P* < 0.05), although blood burden remained detectable until at least 72 h post-infection, which differs from ΔCPS2-infected mice (Fig. 4B and Fig. S8).

Furthermore, plasmatic levels of different pro-inflammatory mediators (12 h post-infection) were evaluated to investigate exacerbated systemic inflammation. The levels were elevated in SS2-infected mice, whereas they were undetectable in ΔCPS2-infected mice (Fig. 4C). Globally, no differences were observed in systemic inflammation between SS2-infected mice and those infected with SS2to7, SS2to9, or SS2to14 (Fig. 4C). However, a significant increase in the production of all the inflammatory mediators was observed in SS2to8-infected mice (*P* < 0.05), in accordance with the results on mortality observed above (Fig. 4A). Meanwhile, plasmatic levels of all mediators were significantly decreased in SS2to4-infected mice compared to SS2 (*P* < 0.05), although levels were detectable (Fig. 4C). Notably, infection with SS2to3 resulted in a significant reduction of most pro-inflammatory mediators compared to SS2, though reduction was not as great as with SS2to4 (Fig. 4C).

### Serotype switching can differentially modulate *S. suis* virulence in piglets

Impact of serotype switching on *S. suis* virulence was subsequently evaluated in the natural host of this bacterium by an experimental intranasal infection model, representing the natural route of exposure to *S. suis*. The mutants were divided into two experiments (experiment I: SS2, ΔCPS2, SS2to4, or SS2to7; experiment II: SS2, SS2to3, SS2to8, or SS2to14) (Table 2). Virulence of the SS2to9 was not evaluated for ethical reasons, since no differences were observed in host cell interactions assays *in vitro* nor in the mouse infection model. In experiment I, none of the ΔCPS2-infected pigs developed any clinical signs of infection, while all SS2-infected pigs showed clinical signs of systemic and/or central nervous system infection, including lame and shivering (Table S3). In fact, three out of four SS2-infected pigs were euthanized at 3 or 4 days post-infection (dpi) due to severity of clinical signs (Table 2 and Table S3). The inoculated strain was recovered from the blood and several organs, including the joints and brain, in all SS2-infected pigs (Table 3 and Table S4). Recovery of SS2 from the joints and brain was also confirmed in the animals presenting lameness or shivering (Table 3 and Table S4). Meanwhile, recovery of the inoculum was not observed from any of the investigated sites in the ΔCPS2-infected pigs, except for the tonsils (two pigs) and the liver (one pig) (Table 3 and Table S4). All SS2to4- and three of SS2to7-infected pigs presented no clinical signs of infection (Table 2 and Table S3), which were, except for the tonsils and a single organ, negative for bacterial recovery (Table 3 and Table S4). However, one of the SS2to7-infected pigs developed shivering, and bacteria were only recovered from the brain and tonsils (Table S4).

**Table 2.** *S. suis* swine infection outcomes and clinical diseases

| Exp.<br>no.-<br>Group<br>no. | Strain | Infectio<br>n dose<br>(CFU) | Mortality<br><sup>a</sup> | Morbidity<sup>b</sup> | Body<br>temp<br>>40.5°C | Description of clinical signs |
| --- | --- | --- | --- | --- | --- | --- |
| I-1 | SS2 | 2.0 ×<br>10<sup>9</sup> | 1/4 | 4/4 | 4/4 | Lameness (3/4)<br>Symptoms improved in one of the<br>pigs<br>Shivering with vomition (1/4) |
| I-2 | ΔCPS2 | 2.9 ×<br>10<sup>9</sup> | 0/4 | 0/4 | 0/4 |  |
| I-3 | SS2to4 | 2.8 ×<br>10<sup>9</sup> | 0/4 | 0/4 | 0/4 |  |
| I-4 | SS2to7 | 3.1 ×<br>10<sup>9</sup> | 1/4 | 1/4 | 1/4 | Shivering and clearly uncoordinated |
| II-1 | SS2 | 1.4 ×<br>10<sup>9</sup> | 0/4 | 0/4 | 2/4 | Slight fever at 4 dpi (2/4).<br>Slight inactivity at 5 dpi (4/4).<br>All animals subsequently recovered |
| II-2 | SS2to3 | 2.8 ×<br>10<sup>9</sup> | 0/4 | 0/4 | 0/4 |  |
| II-4 | SS2to8 | 1.2 ×<br>10<sup>9</sup> | 1/4 | 1/4 | 2/4 | Inactive and lame |
| II-5 | SS2to1<br>4 | 1.2 ×<br>10<sup>9</sup> | 0/5 | 0/5 | 0/5 |  |

<sup>a</sup> Number of pigs to reach predefined clinical end point (see **Text S8** for more detail).
<sup>b</sup> Number of pigs having a score of >1 on attitude or locomotion.
Abbreviations: Exp., Experiment; dpi, days post-infection.

**Table 3.**
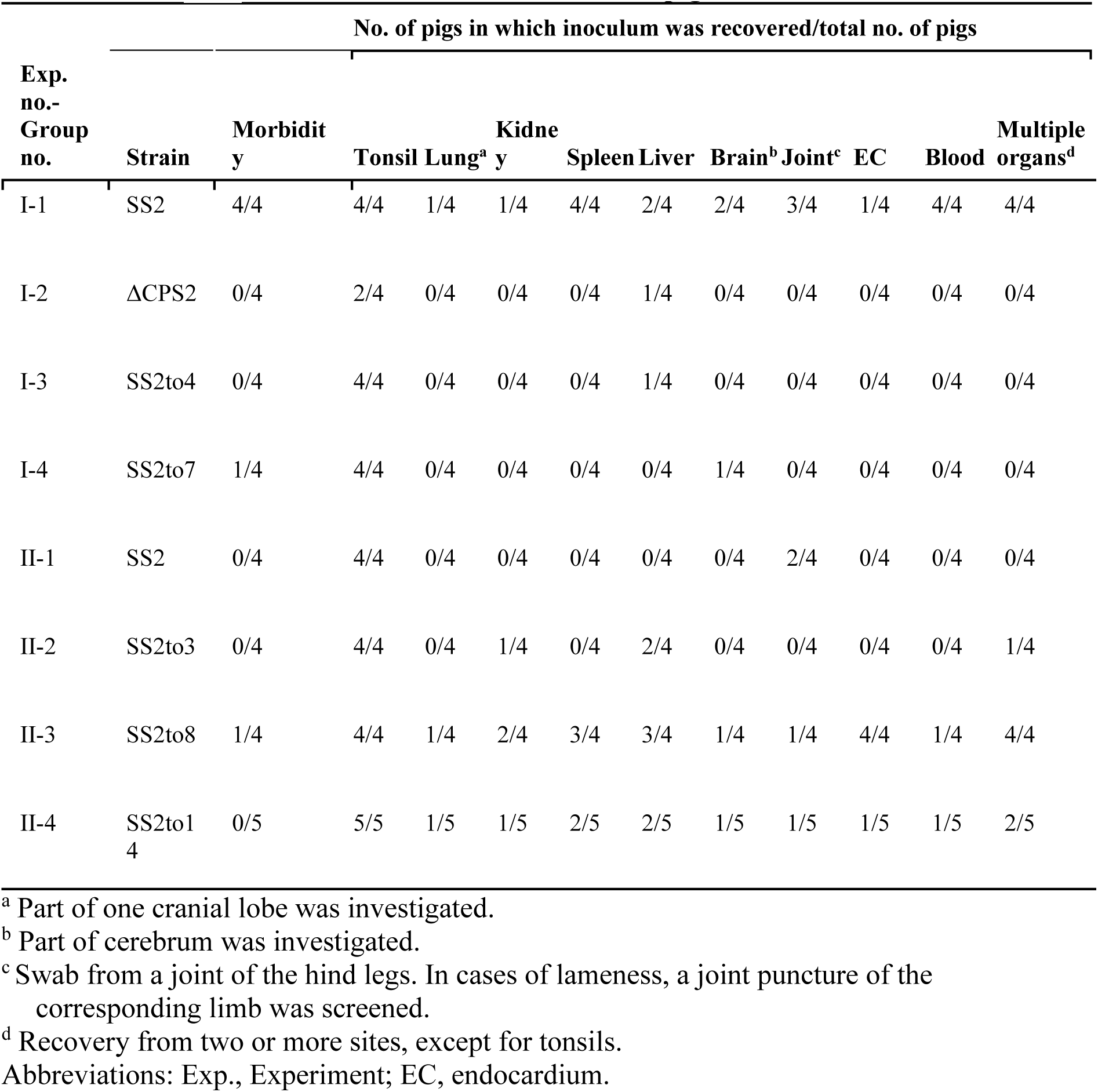
Recovery of inoculated strains from infected piglets

Unfortunately, none of the SS2-infected pigs developed clinical signs in experiment II, with recovery only from the tonsils and joints (Table 3 and Table S4), although slight fever was observed 4 dpi (Table 2 and Table S3). These difference in results of SS2 between experiments may be due to the pigs being used originated from different suppliers. Although most SS2to3-, SS2to8-, or SS2to14-infected pigs showed no clinical signs, one of the SS2to8-infected pigs developed clinical symptoms, including inactivity and clear incoordination (Table 2 and Table S3). Nevertheless, SS2to14 was recovered from the blood and organs of one of the infected pigs. Excluding this individual, however, bacterial recovery was mostly negative for SS2to3- or SS2to14-infected pigs. Meanwhile, bacteria were recovered from multiple organs in all the SS2to8-infected pigs, though recovery from blood was recorded in only the individual presenting clinical symptoms (Table 3 and Table S4).

## Discussion

This study provides the first evidence that serotype switch in *S. suis* can definitively modify the interactions with host cells and *in vivo* (Summarized in Table 4). CPS expression of *S. suis* serotypes 2, 9, and 14 plays critical roles on colonization and anti-phagocytic activity, important steps of the pathogenesis [18, 19, 29]. In this study, under almost the same genetic background of the serotype 2 strain P1/7 (SS2), only switching to serotype 7 or 8 changed the adhesion pattern of SS2 to porcine tracheal epithelial cells. Regarding anti-phagocytic activity, no significant or minor difference was observed by serotype switching. By further evaluation on the effects on serotype-switching using *ex vivo* (blood) and *in vivo* infection models (mouse and pig), only mutants switched to serotype 4 or 8 showed a marked and consistent impact on several bacterial virulence traits. The CPS4 conferred to *S. suis* a non-virulent phenotype characterized by increased susceptibility to killing by mouse and pig blood, reduced bacteremia in mice, diminished cytokine production (*in vitro* and *in vivo*), and low bacterial recovery from internal organs in pigs. In marked contrast, the CPS8 conferred to *S. suis* an hyper-virulent phenotype characterized by high capacity to multiply in pig blood, high bacteremia (mice) and organ dissemination (pigs), and increased capacity to induce a cytokine storm (*in vitro* and *in vivo* in the mouse model). It should be noted that switching to serotype 14 or 9 (variant) had no major effects on *S. suis* virulence or its interactions with the host either *in vitro* or *in vivo* in the mouse model. Meanwhile, serotype switch to CPS7 or CPS3 has restricted impact and affected few of the evaluated parameters. The SS2to7 mutant has slightly increased susceptibility to killing by pig blood and reduced virulence in the swine infection model, being mainly recovered from tonsils. The SS2to3 mutant presented increased susceptibility to killing by mouse and pig blood, slightly reduced bacteremia in mice, and diminished capacity to induce cytokine production *in vivo*. Though serotype 3 CPS expression still caused *S. suis*-induced host death, clinical signs were less severe than those caused by SS2 in the mouse model. None of the pigs infected with SS2 developed clinical signs in experiment II, so a reduced virulence of SS2to3 mutant could not be definitively confirmed in the natural host. Overall, results obtained with the different mutants confirmed the delicate balance between bacterial burden, systemic dissemination, level of the inflammatory response, and clinical outcome [28, 30, 31]. Given that only different CPSs were expressed between mutants, these differences in effects depending on switched serotypes might be due to differential cell wall component exposure, including adhesins and immunostimulatory components, and/or recognition of certain motifs of specific *S. suis* CPSs by unknown host cell receptors.

**Table 4.**
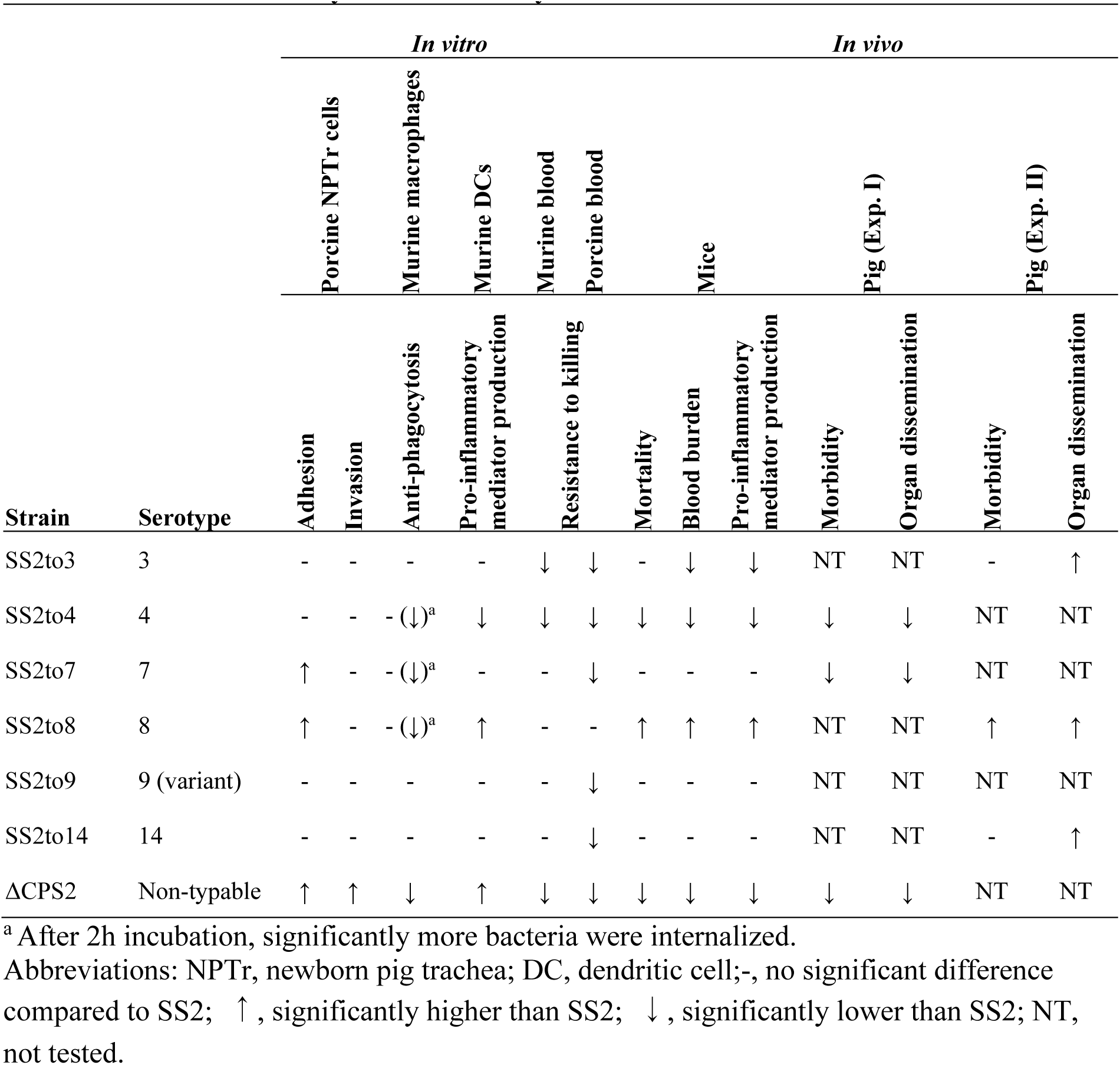
Summary of the effects caused by serotype switching from serotype 2 on *in vivo* and *in vitro* virulence analyzed in this study

This work also highlighted the complexity of *S. suis* host-pathogen interactions and the carefulness required when analyzing data from single cell type cultures *vs.* more complex biological systems (such as blood). For instance, neutrophils and monocytes are the main phagocytes in blood, with little to no macrophages being present. Therefore, results obtained with macrophages might not necessary reflect *S. suis* fitness in blood, but rather mimic the situation in tissues. Similarly, the interactions of *S. suis* with swine blood leukocytes are more complex than those evaluated when using mouse blood due to the presence of swine antibodies reacting against the bacteria. Thus, by using multiple *in vitro* and *in vivo* models, a more comprehensive analysis is obtained.

In *Streptococcus pneumoniae*, strict evaluations of the CPS effects using CPS switch mutants have already been performed, and several studies demonstrated that capsule type affected resistance to both complement C3b deposition and opsophagocytic uptake [32], nonopsonic neutrophil-mediated killing [33], and adhesion to the pharyngeal or lung epithelial cells [34]. Some of these studies also indicated the effect on virulence within the respiratory tract [34], colonization [33], survival in blood [32], and brain injury [33] by *in vivo* infection models. The structure and composition of CPS8 of *S. suis* is known to be identical to that of *S. pneumoniae* serotype 19F [13], with serotype 19F pneumococcus mutant being shown to be the most resistant to non-opsonic killing by human neutrophils among the mutants [33], suggesting that this structure of CPS provides the bacteria with high resistance to killing in blood. Previous studies using serotype-switched mutants [33, 36] also showed that CPS type affects the degree of encapsulation and growth phenotype due to the difference in metabolic costs for producing capsule between CPS types. These points should be evaluated in *S. suis* in the future.

In conclusion, these data demonstrate that serotype switching in *S. suis* serotype 2 can modulate host cell interactions and virulence. Among the tested serotypes, switch to serotype 8 increased the virulence. Although it remains unknown whether *S. suis* serotype switching affects virulence in humans, one serotype 8 strain having a genetic background similar to virulent serotype 2 clinical isolates has already been recovered (unknown source: pubMLST: http://pubmlst.org/ssuis/). Therefore, these results clearly demonstrate that more attention should be given to serotype switching in *S. suis* with regards to both commensal and pathogenic strains.

## Materials and methods

### Ethics statement

The animal experiments in this study were approved by the institutional committees for Ethics of Animal Experiments of the National Institute of Animal Health Japan (approval numbers 17-002, 17-010, and 17-085) and by the Animal Welfare Committee of the University of Montreal (approval number Rech-1570). Both committees formulated the guidelines and policies required to meet and adhere to the standards in the Guide for the Care and Use of Laboratory Animals.

### *S. suis* culturing

The *S. suis* strains used in this study are listed in Table 1. The serotype 2 strain P1/7 (SS2 in this study) [20] was used as the parental strain for construction of the serotype-switched mutants. P1/7 belongs to CC1 and was shown to be induced to a competent state using XIP [17]. *S. suis* strains of serotypes 3, 4, 7, 8, 9, and 14 were used as donors to construct the serotype-switched mutants. All strains were cultured overnight on Todd-Hewitt (TH) agar (Becton Dickinson, Franklin Lakes, NJ, USA) at 37°C with 5% CO2 unless indicated otherwise. Chloramphenicol was added to the medium at 5 μg/mL, when needed.

### General molecular biology techniques

All PCRs were completed using the iProof HF Master Mix (BioRad Laboratories, Hercules, CA, USA) and QIAGEN Multiplex Master PCR Mix (Qiagen, Hilden, Germany) according to the manufacturers’ instructions. The PCR primers used in this study are listed in Table S5. The amplified PCR products were purified using the QIAQuick PCR Purification Kit (Qiagen) and sequenced on a 3130*xl* Genetic Analyzer (Applied Biosystems, Foster City, CA, USA) using a BigDye Terminator v3.1 Cycle Sequencing Kit (Applied Biosystems) where required. The sequence assembly of the PCR products was performed using SEQUENCHER 5.4 (Gene Codes Corp., Ann Arbor, MI, USA).

### Construction of serotype-switched mutants and non-encapsulated mutant

An outline of the approach developed for the construction of the serotype-switched mutants is represented in Fig. S1. First, a non-encapsulated mutant whose *cps* locus was replaced with a chloramphenicol resistance gene (ΔCPS2tocat) was generated from SS2. Then, the ΔCPS2tocat was transformed with whole genome of donor strains to yield the desired serotype-switched mutants through the replacement of the *cat* with the donor *cps* locus (See Text S3 for more detail). For generation of the markerless non-encapsulated mutant, blue-white screening method using 5-bromo-4-chloro-3-indoxyl-α-L-fucopyranoside (X-α-L-fucopyranoside) was performed as represented in Fig. S2 (See Text S4 for more detail).

### *S. suis* growth measurements

Strains were streaked onto TH agar plates and incubated overnight at 37°C with 5% CO2 and then subcultured in TH broth to an optical density 600 nm (OD600) of 0.6 using a spectrophotometer Ultrospec 2100 (Biochrom Ltd., Cambridge, UK). After adding 1/500 of the volume of each adjusted culture diluted 1,000 times by TH broth to TH broth, the cultures were incubated at 37°C under air plus 5% CO2 conditions. The CFU(/mL) of each of the cultures was measured at 2, 4, 6, 8, 10, 12, and 14 h after incubation by plating serial dilutions on TH agar.

### Confirmation of serotype switching

Serotyping, cell surface hydrophobicity test, TEM, measurement of CPS yields, NMR spectroscopy were performed to confirm well-encapsulation and serotype switching as previously described [27, 37, serotyping and TEM; 38, hydrophobicity tests; 9,11,12,13, CPS purification and NMR] (see Text S5 for more detail).

### Whole genome sequence analyses

Whole genome draft sequences were determined using Illumina HiSeq X ten sequencing platform at the Beijing Genomics Institute (Shenzhen, China) or Illumina NovaSeq platform at Novogene Corporation (San Diego, CA, USA) (See Text S6 for more detail). The final draft genome sequence of each of the mutants was then mapped and aligned with the publicly available complete genome sequence of strain P1/7 using Geneious Prime ver. 2019.1.1 (Tomy Digital Biology, Tokyo, Japan) with the default parameters.

### In *vitro* assays for evaluation of impacts on serotype switching

Adhesion and invasion assays using the porcine tracheal epithelial NPTr cell line, phagocytosis assays using J774A.1 murine macrophages, murine whole blood bactericidal assay using blood collected from 6- to 10-week-old C57BL/6J mice and from a five-week-old piglet, and measurement of pro-inflammatory mediator production by DCs generated using the femur and tibia of C57BL/6J mice were performed as previously described [19, 28, 39]. (see Text S7 for more detail).

### *In vivo* assays for evaluation of impacts on serotype switching

Mouse infections were performed using 10-12 six-week-old male and female C57BL/6J mice per group via intraperitoneal inoculation (dose of 1 × 10^7^ CFU/mouse) for survival and blood bacterial burden evaluation as previously described [28]. Plasma (systemic) pro-inflammatory mediators were measured using blood collected from eight mice intraperitoneally infected with 1 × 10^7^ CFU 12 h post-infection as previously described [28]. Pig infections were performed for evaluation of appearance of symptoms and organ dissemination using 4-5 five-week-old crossbred male and female piglets per group purchased from Shokukanken Inc. (Gunma, Japan) or CIMCO Co. Ltd. (Tokyo, Japan). Infections were carried out via intranasal inoculation (dose of 2 × 10^9^ CFU) for survival as previously described [40] and divided into two experiments per four groups (Experiment I: SS2, ΔCPS2, SS2to4, and SS2to7; experiment II: SS2, SS2to3, SS2to8, and SS2to14) (see Text S8 for more detail).

### Statistical analyses

Normality of data distribution was verified using the Shapiro-Wilk test and Mann-Whitney rank sum tests were performed to evaluate statistical differences between groups. Data are presented as mean ± SEM or as geometric mean. Log-rank (Mantel-Cox) tests were used to compare survival between groups of mice. *P* < 0.05 was considered statistically significant.

### Data availability

All sequences determined in this study were deposited in the DDBJ/ENA/GenBank databases under the accession numbers (P1/7, WABV00000000; ΔCPS2tocat, WABW00000000; SS2to3, WABX00000000; SS2to4, WABY00000000; SS2to7, WABZ00000000; SS2to8, WACA00000000; SS2to9, JABMDA000000000; SS2to14, WACB00000000; MO690, WACC00000000; MO691, WACD00000000; MO941, WACE00000000).

## Acknowledgments

This work was funded by the JSPS KAKENHI grants #18H02658 (MO and TS) and #26870840 (MO), as well as by the Natural Sciences and Engineering Research Council of Canada (NSERC) grants #04435 (MG) and #342150 (MS). JPA and GGD are recipients of an Alexander Graham Bell Graduate Scholarship – Doctoral Program from NSERC. MS is the holder of a Canada Research Chair – Tier 1. The funders had no role in study design, data collection and interpretation, or the decision to submit the work for publication.

The authors would like to thank Sonia Lacouture for technical help and advice, Mariane Grzebyk and Claudia Duquette for technical assistance with the production and purification of the CPSs, Kaori Tosaki, Kennosuke Sugie, Koujiro Yoshizaki, Yusuke Abeto, and Hirotaka Itoh for TEM analysis, and Han Zheng for providing information on genome sequence of serotype 9 strains. Computational resources were partly provided by the Data Integration and Analysis Facility, National Institute for Basic Biology, Japan.

The authors would like to thank Sonia Lacouture for technical help and advice, Mariane Grzebyk and Claudia Duquette for technical assistance with the production and purification of the CPSs, Kaori Tosaki, Kennosuke Sugie, Koujiro Yoshizaki, Yusuke Abeto, and Hirotaka Itoh for TEM analysis, and Han Zheng for providing information on genome sequence of serotype 9 strains. Computational resources were partly provided by the Data Integration and Analysis Facility, National Institute for Basic Biology, Japan.

## Supporting information

**Fig S1. Reported composition and structure of the *S. suis* serotype 2, 3, 7, 8, 9, and 14 CPSs.** Monosaccharide symbols follow the Symbol Nomenclature for Glycans System (Varki A, Cummings RD, Aebi M, Packer NH, Seeberger PH, Esko JD, et al. Symbol nomenclature for graphical representations of glycans Glycobiology. 2015;25: 1323-1324). Abbreviations: D-6d-*xyl*HexNAc, 2-acetamido-2,6-dideoxy-D-*xylo*-hexose; 4NAc, 4-acetamido; 4N, 4-amino.

**Fig S2. Diagram of the procedure used for construction of the *S. suis* serotype-switched mutants.** The procedure consists of five steps. Construction of the non-encapsulated mutant (step ***1***) is the most important step of the procedure, as it is essential for the following screening and selection steps. Due to a lower buoyancy density of encapsulated bacterial cells than those of the non-encapsulated cells, a density gradient centrifugation with Percoll (step ***3***) was used to screen encapsulated (i.e., serotype-switched) transformants from ΔCPS2tocat transformed with genome DNA of a donor strain in step ***2***. Moreover, encapsulated transformants were further selected by the differences in how the precipitations were formed in a static liquid medium (step *4*) due to the elevated hydrophobicity of non-encapsulated cells. Abbreviation: CP, chroramphenicol; XIP, *sigX*-inducing peptide; TH, Todd-Hewitt.

**Fig S3. Diagram of the procedure used for construction of the markerless *S. suis* non-encapsulated mutant.** Abbreviations: CP, chroramphenicol; XIP, *sigX*-inducing peptide; TH, Todd-Hewitt.

**Fig S4. Growth curves of the different *S. suis* serotype-switched mutants.** Growth curves of P1/7, non-encapsulated mutant (ΔCPS2) and serotype-switched mutants (SS2to3, SS2to4, SS2to7, SS2to8, SS2to9, and SS2to14) derived from P1/7 are shown.

**Fig S5. 500 MHz ^1^H NMR spectra of the *S. suis* serotype-switched mutant CPSs. (A)** SS2to3, resonance reporter signals (85°C): δ 4.56 and 4.33 (anomeric), 2.01 and 1.94 (acetyl methyl), and 1.22 (6-deoxy sugar methyl); **(B)** SS2to7, resonance reporter signals (25°C): δ 5.68, 5.43, 5.07, and 4.58 (anomeric), 2.02 (acetyl methyl), as well as 1.33 and 1.24 (6-deoxy sugar methyl); **(C)** SS2to8, resonance reporter signals (25°C): δ 5.49, 5.01, and 4.91 (anomeric), 2.08 (acetyl methyl), and 1.30 (6-deoxy sugar methyl); **(D)** SS2to9, resonance reporter signals (50°C): δ 5.44, 5.40, 5.00, 4.99, 4.96, 4.80, 4.76, and 4.72 (anomeric), 2.04 (acetyl methyl), as well as 1.27, 1.27, and 1.24 (6-deoxy sugar methyl); **(E)** SS2to14, resonance reporter signals (77°C): δ 4.77, 4.62, 4.50, 4.50, and 4.45 (anomeric), 2.05 and 2.03 (acetyl methyl), 2.68 (Neu5Ac H-3e), and 1.68 (Neu5Ac H-3a). Except for SS2to9 CPS, the slight differences in chemical shifts compared to published values [references 9, 12, and 13 in the text] can be attributed to different sample concentration and pH, internal reference, and spectral acquisition temperature. Two-dimensional (2D) correlation spectroscopy (COSY) experiments, and additionally for SS2to9 CPS the 2D heteronuclear single-quantum coherence (HSQC) experiment, were also performed, and the observed cross-peaks were in complete support of the structures. Abbreviation: Sug, 6d*xyl*HexNAc-4-ulo.

**Fig S6. Difference in CPS between SS2to9 and strain 1273590.** (A) Composition and structure of CPSs. (B) Nucleotide sequence alignment between the *cps* loci. Each schematic representation shows the analysis data using Geneious Prime. Below the bottom panel are displayed the descriptions for each color of the different drawings. Nucleotide sequence of *cps* locus of 1273590 was extracted from its draft genome sequence (Accession no. SRS1751390). Glycosyl transferase genes, *cps9F, cps9G, cps9H, cps9I*, and *cps9K* were appended.

**Fig S7. Replacement between *cps* loci. (A)** SS2to3, **(B)** SS2to4, **(C)** SS2to7, **(D)** SS2to8, **(E)** SS2to9, and **(F)** SS2to14. Each schematic representation shows the analysis data using Geneious Prime on the sequence alignment between the *cps* loci and their flanking regions of the serotype-switched mutants and donor strains (upper part) and between the *cps* loci of the serotype-switched mutants and reference serotype strains (lower part). Below the bottom panel are displayed the descriptions for each color of the different drawings.

**Fig S8. Blood bacterial burdens at 48 h and 72 h post-infection of mice inoculated with the different *S. suis* strains and mutants.** Data represent the geometric mean (n = 10-12). A blood bacterial burden of 2 × 10^9^ CFU/mL, corresponding to average burden upon euthanasia, was attributed to euthanized mice. n.d. denotes not determined. An asterisk denotes a significant difference with SS2 by Mann-Whitney rank sum test (p < 0.05).

**Table S1. Purification yields of the CPS from the different *S. suis* strains and serotype-switched mutants**

**Table S2. Mutated genes present in the serotype-switched mutants and their amino acid identities with those corresponding of strain P1/7**

**Table S3. Daily score of individual piglets**

**Table S4. Reisolation of the infection strain from each piglet**

**Table S5. Primers used in this study**

